# Sex-dependent involvement of lateral septum astrocytes in social fear: Role of oxytocin receptor signaling

**DOI:** 10.64898/2026.02.01.703181

**Authors:** Laura Boi, Rohit Menon, Clémence Denis, Kai-Yi Wang, Hugues Petitjean, Barbara Di Benedetto, Alexandre Charlet, Inga D. Neumann

## Abstract

Astrocytes are now widely recognized as important modulators of synaptic plasticity and socio-emotional behaviors. Recent studies highlight their involvement in anxiety- and depressive-like behaviors, particularly via oxytocin (OXT) signaling. While the specific contributions of astrocytes remain largely unexplored, the role of OXT receptor (OXTR) signaling in the lateral septum (LS) in regulating social fear expression has been well characterized. Here, we studied the differential contribution of astrocytic OXTR signaling using a social fear conditioning (SFC^+^) paradigm. We found the highest abundance of astrocytes, and especially of OXTR-expressing (OXTR^+^) astrocytes, within the caudal LC (LSc) compared to the rostral LS in both male and female mice. Interestingly, female mice displayed a significantly higher number of astrocytes and OXTR^+^ astrocytes in the LSc in comparison to males. However, social fear acquisition resulted in dynamic changes in LSc astrocytic morphology and calcium activity in male mice. Furthermore, we showed that pre-SFC acquisition pharmacology-induced loss of local astrocytic function facilitated the extinction of social fear in males. In support, astrocyte-specific OXTR knockdown in the LSc also facilitated social fear extinction in both males and females. Taken together, our study identifies OXTR-signaling in LSc astrocytes as a crucial component in the mechanisms underlying the regulation of social fear in a sex-dependent manner.

## Introduction

Social interactions are essential for the survival and well-being of individuals. However, when responses to social cues become maladaptive, as in social anxiety disorder, they pose a challenge to individual health and a burden for society. Social anxiety disorder, also referred to as social phobia, is characterized by an intense and persistent fear of social situations ^1^. Social anxiety disorder is the 3^rd^ most common anxiety disorder, with a lifetime prevalence of 12 % and a rising global incidence ^2, 3^. Standard treatments combine psycho- and pharmacotherapeutic approaches ^4, 5^, although their efficacy remains limited. Approximately 40% of patients experience treatment resistance and relapse ^6, 7^. These challenges underscore the need for a more comprehensive examination of the underlying mechanisms. A relevant animal model for social anxiety disorder is the operant conditioning-based social fear conditioning (SFC) paradigm that induces robust social avoidance in both male and female mice ^8, 9^.

Oxytocin (OXT), a neuropeptide known for its pro-social and anxiolytic effects, has emerged as a promising therapeutic target for anxiety-related psychopathologies ^10–12^. Our research has implicated OXT signaling specifically within the lateral septum (LS) in attenuating or reversing SFC-induced social fear expression ^9, 13^. Particularly, the heightened activity of OXT signaling in the LS of female mice during lactation ^9^ and in male mice post-ejaculation ^14^ were found to attenuate SFC-induced social fear expression. The LS is a heterogeneous brain structure of the limbic system known to regulate various socio-emotional behaviors ^15, 16^. Neurons of the LS respond dynamically to social fear acquisition and extinction, as reflected in neuronal activity patterns and the expression of genetic and epigenetic markers ^17–19^.

While OXT research in the context of SFC has predominantly focused on neurons, the role of astrocytes slowly emerges. Astrocytes are recognized as key modulators of synaptic plasticity and emotional behaviors ^20–22^. OXT receptors (OXTR) are expressed in both neurons and astrocytes across the brain ^23–26^. Pioneering studies have shown that OXT induces structural changes in hypothalamic astrocytes, consequently affecting neuronal network dynamics ^27^. Specifically, both endogenous OXT released during heightened activity states of the OXT system, such as lactation ^28^, acute stress ^26^ as well as infusion of synthetic OXT ^26, 29, 30^ trigger astroglial process retraction, which enhances neuronal communication. This highlights the impact of OXT on the spatial organization of synapses and neuron-astrocyte interactions ^25, 31, 32^.

Recent evidence indicates that astrocytic OXT signaling plays a significant role in general anxiety-related behaviors ^25, 29, 33–35^. In addition, OXTR has been found in a subset of ∼20% of LS astrocytes of both rats and mice, a similar proportion to that observed in other brain regions. However, the specific role of LS OXTR^+^ astrocytes in regulating social fear remains unknown. In addition, sex appears as a critical variable of social anxiety disorder, which disproportionately affects more women (DSM-5). Whether the observed sex differences in the occurrence of social anxiety disorder are linked to the differential OXTR expression in the brain ^36–38^ and astrocytic functions ^39, 40^ remains unknown. Given the limited treatment options for social anxiety disorder, it is interesting to investigate the role of astrocytic OXT signaling in mediating social fear expression. To this end, we first studied (i) sex differences in social fear expression in the SFC paradigm, (ii) sex differences in the distribution pattern of astrocytes within the LS and the astrocytic response (morphological and Ca^2+^ levels) to social fear acquisition, and (iii) the functional involvement of LS astrocytes in regulating social fear expression during acquisition and extinction. Next, we studied (iv) the distribution of OXTR-expressing (OXTR^+^) astrocytes within the LS and (v) the effects of social fear acquisition on the responsiveness of these OXTR^+^ astrocytes to the selective OXTR agonist TGOT. Finally, (vi) we studied the functional role of LS OXTR^+^ astrocytes in modulating social fear extinction in male and female mice.

## Materials and Methods

### Animals

Wildtype CD1 virgin female and male mice (Charles River, Sulzfeld, Germany, 9-12 weeks of age) were housed under standard laboratory conditions (12/12 h light/dark cycle, 22°C, 60% humidity, food, and water *ad libitum*) in groups of 4 mice after arrival at the animal facility. After surgery, mice were single-housed for either 5 days (for guide cannula implantations) or 3 weeks (for viral microinfusions) before behavioral experiments commenced. Experiments were performed in the light phase between 08:00h and 12:00h. They were approved by the government of Unterfranken, and performed in accordance with EU guideline 2010/63/EU and ARRIVE guidelines (Kilkenny et al., 2010).

### Social fear conditioning (SFC) paradigm

The SFC paradigm was performed as previously described ^8, 9^.

#### Social fear acquisition (day 1)

Female and male mice were individually placed in the conditioning chamber (TSE System GmbH, Bad Homburg, Germany) and allowed to habituate for 30s before being exposed to an empty wire-meshed cage (non-social stimulus), which they could freely investigate. Then, the non-social stimulus was replaced with an identical wire-meshed cage containing a sex- and age-matched conspecific (the social stimulus). Non-conditioned mice (SFC^−^) were allowed to freely investigate the social stimulus for 3 min, while conditioned mice (SFC^+^) received a foot shock (US: Unconditioned Stimulus; mean 2.4-foot shocks/mouse) each time they approached, sniffed, and made physical contact with the social stimulus (CS: Conditioned stimulus). After the first CS-US pairing, we waited an additional 6 min to see whether the mice made further social contact. However, after the 2nd CS-US pairing, the observation period for social contact was reduced to 2 min, after which the test mouse was returned to their home cage.

#### Social fear extinction (day 2)

One day after social fear acquisition, SFC^+^ and SFC^−^ mice were exposed to 3 non-social stimuli (empty cages) followed by consecutive exposure to 6 different same-sex, same-age unfamiliar social stimuli (placed in a small empty cage) in their home cage for 3 min each, with a 3-min inter-exposure interval.

#### Social fear extinction recall (day 3)

One day after social fear extinction, mice were consecutively exposed to 6 different same-sex, same-age unfamiliar social stimuli (placed in a small empty cage) in their home cage for 3 min, with a 3-min inter-exposure interval.

During social fear extinction and recall, two key behaviors were analyzed: the investigation time, defined as the time the mouse spent interacting and in direct contact with the non-social or social stimulus ^8^, and the vigilance time, defined as the time spent moving or stationary (e.g. behaviors such as freezing and stretch-attend posture) with the head oriented toward the stimulus ^41^. Animal behavior was recorded and analyzed by a trained observer, blind to the treatment conditions, using the JWatcher program (V 1.0, Macquarie University and UCLA).

### Stereotaxic implantation of guide cannulae and bilateral drug infusion into the LS

For bilateral drug infusions, guide cannulas (8 mm long, 23 G, Injecta Gmbh, Germany) were stereotaxically implanted 2 mm above the right and left LSc (+0.15 mm, ± 0.5 mm, -1.6 mm from bregma) ^42^ under isoflurane anesthesia (Forene, Abbott, Wiesbaden, Germany) under semi-sterile conditions ^8, 9, 13^. The guide cannulas were secured in place by dental cement (Kallocryl, Speiko-Dr. Speier Gmbh, Germany) and closed with a stainless-steel stylet (27 G). After surgery, mice were single-housed for 5 days and handled daily prior behavioral testing to reduce unspecific stress responses during the experiment.

Mice received bilateral LS infusions of either the astrocytic inhibitor L-AAA (25 µg/µl; 0.2 µl/hemisphere) or vehicle (PBS, 10x, 0.2 µl/hemisphere; 27 G infusion cannula, 10 mm long) 3 days before social fear acquisition to allow efficient astrocytic inhibition. The dose and timing of L-AAA were selected based on previous studies ^43, 44^. The correct infusion sites were histologically verified and, accordingly, animals with misplaced guide-cannula were removed from the statistical analysis. To verify the effectiveness of L-AAA in inhibiting astrocytes, two different dosages of L-AAA (10/25 µg/µl) were infused bilaterally into the LSc. Immunohistochemical and electrophysiological analyses were performed to measure GFAP fluorescence intensity in a 600-µm square region of interest, positioned adjacent to the infusion site to account for potential tissue damage (Fig. S1A-C).

### Stereotaxic virus microinfusions into the LSc

To induce astrocyte-specific knockdown of OXTR mRNA within the LSc, where we identified a higher percentage of OXTR^+^ astrocytes, an adeno-associated virus (AAV) expressed under the GFAP promoter, combined with short hairpin RNA (shRNA) and mCherry fluoroprotein was used (AAV6-GFAP-OXTR-mCherry-shRNA constructs; OXTR-shRNA; 3.28×10^13^ GC/ml, VectorBuilder). Mice received stereotaxic microinfusions (280 nl / hemisphere) of either the OXTR-shRNA vector or a control vector (scrambled RNA oligonucleotide; scrRNA) via a pulled electrophysiology glass-pipette into the left and right LSc (AP +0.15 mm, ML ±0.5 mm, DV -3.2 mm) ^42^ 21 days before behavioral testing.

### Immunohistochemistry and morphological analysis of astrocytes in the LS

To study the distribution and density of astrocytes within the LS, and the effects of SFC on astrocytic morphology in the LSc, male and female mice were transcardially perfused with paraformaldehyde (PFA) under deep anesthesia 90 min after social fear acquisition. Brains were removed and post-fixed in 4% PFA for 24 h, followed by cryo-protection in 30% sucrose for 2 days and snap-freezing in isopentane. Forty-µm coronal cryocut slices (CM3000, Leica, Germany) containing the LS (Bregma from +1.18 to - 0.15) were kept in PBS containing 5% normal goat serum (Vector Laboratories) and 0.5% TritonX-100 (1 h at room temperature (RT), and, afterwards, incubated with primary antibody against glial fibrillary acidic protein (GFAP) (cs123895; Cell Signaling Technology) and S100ß (S2532, Millipore Sigma) to guarantee a broader coverage of total astrocytes (48 h, 4°C). Appropriate secondary antibody (Anti-rabbit IgG, 7074S; Cell Signaling Technology) was applied (2 h, RT) in the dark. Slides were covered with SuperFrost slides using Roti® Mount FluorCare DAPI (Carl Roth, Germany). Analysis of density was performed by manually counting the number of GFAP/S100β^+^ cells within the rostral (LSr), caudal (LSc) LS.

Because of the higher density of OXTR^+^ astrocytes in the LSc compared with the LSr (Fig. 4C), astrocytic morphology was analysed in the LSc. Images were captured using a confocal laser-scanning microscope (CLSM; Zeiss LSM 980 Airyscan, Germany). Three z-sections (each 30-µm thick, with a 0.5-µm/z section interval, acquired at 1024×1024 resolution using a 63x objective) were taken per LS subregion. The length of the longest astrocytic process and domain area of each astrocyte were analyzed (ImageJ software, Version 1.53c) ^45^. Results were regularly cross-validated by an independent investigator blinded to the treatment.

### RNAscope and colocalization of OXTR mRNA and GFAP/S100β

To compare the number of OXTR^+^ astrocytes along the LS of naïve male and female mice, RNA scope was performed on 20-µm thick coronal slices (Bregma +1.18 to -0.10) using the RNAscope Multiplex Fluorescent V2 kit (Advanced Cell Diagnostic, Acdbio) ^24^. Slices were processed using a probe for OXTR (483671-C3) and the TSA Plus Fluorophore Cy3 (1:1000; FP1168, Perkin Elmer). To visualize OXTR^+^ astrocytes, RNAscope slices were stained for GFAP and S100β and analysed using a confocal laser-scanning microscope (CLSM, Zeiss LSM 980 Airyscan, Germany). The number of OXTR^+^/ GFAP/S100β^+^ cells was manually counted within the LSr and LSc. Total number of GFAP/S100ß^+^ cells equals 100% (Fig 4C).

### Gene expression analysis

To assess the efficiency of the OXTR knockdown using shRNA, relative OXTR mRNA expression in the caudal septum was measured in both shRNA and scrRNA groups. For details, see Supplementary.

#### RNA isolation

Brain punches from the septum were collected and RNA was isolated with 500 µl of peqGold® TriFast (peqLab, Erlangen, Germany) and mixed with 100 µl of chloroform. After centrifugation (20 min, 17,000 g, 4 °C), the supernatant containing the RNA was precipitated with approximately 200 µl of isopropanol (45% v/v) and stored at -20 °C overnight. The RNA pellet was washed twice with 500 µl of 80% ethanol, centrifuged (15 min, 17,000 g, 4 °C), air-dried for 5–10 min, and resuspended in 7 µl of nuclease-free water. Samples were heated at 70 °C for 5 min while shaking. RNA concentration and purity were measured at 260/280 nm and 230/260 nm, respectively, using a NanoDrop spectrophotometer (ThermoScientific, Waltham, USA).

#### Reverse transcriptase PCR, endpoint PCR, and quantitative PCR (qPCR)

For cDNA synthesis, 300 ng of RNA was adjusted to a final volume of 20 µl with RNase-free water and mixed with 4 µl of UltraScript Butter 5x buffer and 1 µl of Random Hexamers (5 µM final concentration, PCR Biosystem Inc., USA). Negative controls (-RT) contained water instead of the reverse transcriptase to check for DNA contamination. After primer annealing (2 min at 70 °C), 1 µl of each reaction mix was added to the -RT control tubes, and 1 µl of UltraScript 2.0 enzyme (UltraScript 2.0 cDNA Synthesis Kit, PCR Biosystem Inc., USA) was added to the remaining samples. All samples were incubated at 25 °C, 50 °C, 80 °C, and 95 °C for 10 min each using a Mastercycler® nexus X2 (Eppendorf, Wesseling-Berzdorf, Germany). cDNA was stored at -20 °C until further analysis.

### Ex vivo calcium imaging

#### Perfusion, slice preparation and dye loading

In order to reveal the effects of social fear acquisition on the responsiveness of OXTR^+^ astrocytes of the LS to OXT, ex vivo calcium imaging on slices containing the LS was performed as previously described ^25^. Ten min after social fear acquisition, mice were anesthetized with ketamine (Imalgene, i.p., 100 mg/kg), and additionally treated with xylazine (Rompun, i.p., 20 mg/kg) and lidocaine (s.c., 10 mg/kg). Brains were extracted after intracardiac perfusion into ice-cold N-methyl-d-glucamine (NMDG)-aCSF composed of (in mM): NMDG (93), KCl (2.5), NaH2PO4 (1.25), NaHCO3 (30), MgSO4 (10), CaCl2 (0.5), HEPES (20), D-glucose (25), L-ascorbic acid (5), thiourea (2), sodium pyruvate (3), N-acetyl-L-cysteine (10) and kynurenic acid (2), as previously described ^46^. The pH was adjusted to 7.4 using either NaOH or HCl, after bubbling with carbogen (95% O2/5% CO2), which was maintained throughout the experiment.

After brain extraction, 350-μm horizontal slices containing the LS were obtained using a Leica VT1000S vibratome. Next, brain slices were hemisected and placed, for a minimum period of 1 h, before any experiments, in a holding chamber at room temperature containing normal aCSFs. The aCSF is composed of (in mM): NaCl (124), KCl (2.5), NaH2PO4 (1.25), NaHCO3 (26), MgSO4 (2), CaCl2 (2), D-glucose (15), adjusted for pH values of 7.4 with HCL or NaOH, and continuously bubbled in 95% O2/5% CO2 gas. Both NMDG-based and normal aCSF were checked for osmolality and kept between 290 and 310mOsm L−1, bubbled with carbogen (95% O2/5% CO2 gas). Slices were then transferred from the holding chamber to an immersion recording chamber and superfused at a rate of 2 ml min−1 with normal aCSF.

#### Calcium imaging and identification of astrocytes

To identify astrocytes, slices were incubated for 20 min at 35 °C in a sulforhodamine 101 (SR101; 1 μM), a fluorophore labelling astrocytes, containing aCSF. Thereafter, the synthetic calcium indicator OGB1-AM was bulk loaded as previously described ^47^, reaching final concentrations of 0.0025% (20μM) for OGB1-AM, 0.002% for Cremophor EL, 0.01% for Pluronic F-127, and 0.5% for DMSO in aCSF and incubated for 1h at 37°C Slices were then washed in aCSF for at least 1h before imaging. Only astrocytes co-labeled for SR101 and OGB1 were used. The spinning-disk confocal microscope for calcium imaging was composed of a Zeiss Axio Examiner microscope with a ×40 water-immersion objective (numerical aperture of 1.0), mounted with an X-Light Confocal Unit–CRESTOPT spinning disk.

The region of interest was identified using a 4x objective and visualized with the uEye cockpit software. To identify SR101/OGB1-labeled astrocytes, the MetaFluor software was used, and the following parameters were applied to illuminate the cells: 20ms at 575nm for SR101 and 80ms at 475ms for OGB1 (2Hz, 500ms interval between acquisitions with a spinning disk pinhole diameter and rotation speed of 70 µm and 15.000 rpm, respectively). Images were acquired at 2Hz with an optiMOS sCMOS camera (Qimaging). Cells within a confocal plane were illuminated for 20ms at 575nm for SR101 and 80ms at 475nm for OGB1 using a Spectra 7 LUMENCOR. The different hardware elements were synchronized using MetaFluor 7.8.8.0 (Molecular Devices). Because astrocytes are mechanosensitive ^48, 49^, OXTR agonist TGOT (500nM) was bath-applied for 20s and not puff-applied to avoid mechanical stimulation. All calcium imaging experiments were conducted at room temperature, and cells with an unstable baseline were discarded.

#### Calcium imaging data analysis

All analyses were conducted as described ^47^. Astrocytic calcium levels within the LS were measured in manually outlined regions of interest (ROI) comprising the cell body using ImageJ software. Subsequent offline data analysis was performed using a custom-written Python-based script. To take into account slice micro-movements, the SR101 fluorescence values were subtracted from the OGB1 ones. Then, a linear regression was applied to each trace to correct for photobleaching. Calcium transients were detected using the find_peaks function of the SciPy library as fluorescence variation exceeding 8 time the standard deviation and a prominence exceeding it 5 times. The number of peaks and the area under the curve was quantified before and after the drug application. All data were normalized according to the duration of the recording, and astrocytes were labeled as ‘responsive’ when their AUC or their calcium transient frequency at least doubled after drug application. Because the time after stimulation is longer than the baseline (10min vs. 3.20min), the probability of observing a spontaneous calcium peak is stronger after stimulation. To avoid this bias, astrocytes with only one calcium peak during the whole recording were not considered responsive. Each calcium transient was then isolated, its area was estimated with the trapezoid method, and its duration was measured as its half-maximum full width (HMFW). The rise and decay phases of Ca^2+^ transients were fitted with linear functions and the coefficient of these slopes was used as rise and decay constants. Fiji software was also used on SR101/OGB1 pictures to produce illustrative pictures. All calcium imaging experiments were conducted at controlled room temperature (26°C), and cells with an unstable baseline were discarded.

### Statistical analysis

All data were analyzed using SPSS (Version 26.0, IBM Corp., USA) or GraphPad Prism (Version 8, GraphPad Software, San Diego, USA). All datasets were first tested for normal distribution using SPSS (Version 26.0, IBM Corp., USA). Accordingly, parametric paired and unpaired t-tests were performed between two groups, or non-parametric Wilcoxon and Mann-Whitney U-tests were used when the data did not meet the normal distribution requirements. Analysis of variance (ANOVA) mixed model ANOVA was used when assessing two factors over two or more time points (factor sex, factor treatment x SFC x time, in SFC extinction and recall). Kruskall-Wallis or Geisser-Greenhouse correction was applied when sphericity was violated. Statistical significance was accepted at p ≤ 0.05, while a trend was recognized at p ≤ 0.07. Data is represented as the mean ± standard error of the mean (SEM). For details, see Tables T1 – T15.

## Results

### SFC^+^ female mice exhibit reduced social fear during social fear extinction

The regulation of rodent social behavior is highly sex-dependent ^40, 50–52^. While we have successfully established the SFC paradigm in both male and female mice ^8, 9^, here we directly compared the behavioural responses of the two sexes (Fig. 1A-D) and used vigilance behavior as an additional parameter for assessing social fear expression ^41^. Male and female SFC^+^ mice required a similar number of CS-US pairings to induce social avoidance (Fig. 1B) during the acquisition of social fear (day 1). Prior to extinction training (day 2), SFC^+^ mice of both sexes displayed similar investigation and vigilance times of the three non-social stimuli, indicative of comparable general anxiety levels (Fig. 1C & E, left). Also, during social fear extinction training, SFC^+^ and SFC^−^ mice differed in social investigation times in both males and females (SFC x sex x time F_3.06,61.2_ = 2.907, p = 0.041; Fig 1C). Specifically, during exposure to the first social stimulus (s1), both male (p < 0.001) and female (p = 0.041) SFC^+^ mice displayed profound social fear reflected by reduced investigation times compared to respective non-conditioned mice (SFC^−^), which increased over the extinction period (Fig. 1C right). However, while male SFC^+^ mice showed lower social investigation times of social stimuli 1 and 2 (s1 p < 0.001; s2 p = 0.031 vs. male SFC^−^), female SFC^+^ mice showed lower investigation only of stimulus 1 (p = 0.014 vs. female SFC^−^), indicating faster extinction of social fear. In support, male SFC^+^ mice showed higher vigilance times both during s1 and s2 (p < 0.001 and p = 0.026 vs. male SFC^−^ ; Fig. 1E right), whereas females displayed higher vigilance times only during exposure to social stimuli 1 (p = 0.019 vs. female SFC^−^ ; Fig 1E right), providing further support for faster extinction in female SFC^+^ mice. During recall, we found no group difference in either social investigation or vigilance behavior (Fig. 1D, F).

**Figure 1.**
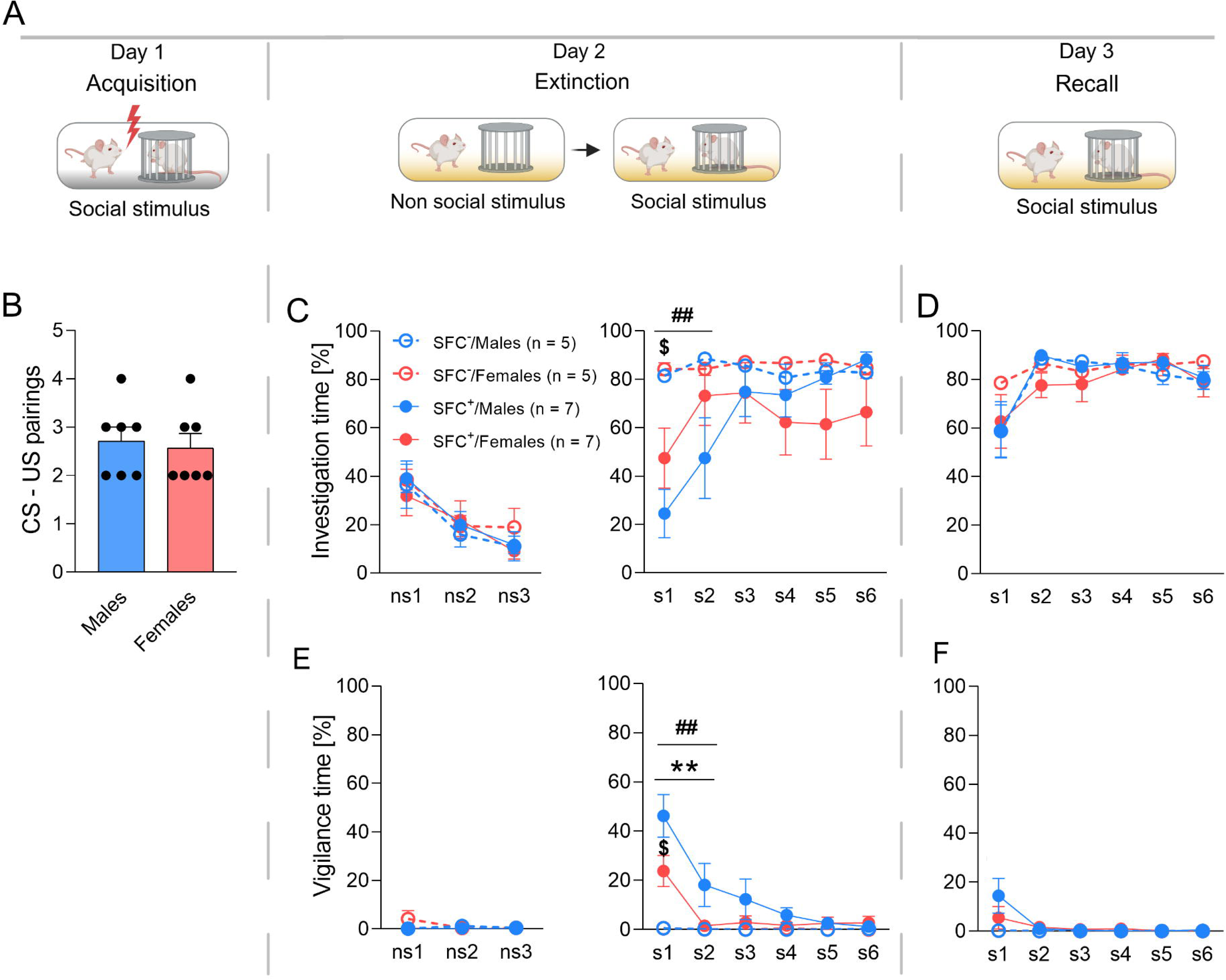
Behavior of male and female mice during social fear acquisition (B), social fear extinction (C, E), and recall (D, F). **A.** Schematic presentation of the experimental protocol. **B.** Number of CS-US pairings necessary to induce social fear in conditioned mice (SFC^+^) during social fear acquisition (day 1). **C.** Percentage of investigation time of the 3 non-social (ns1 - ns3; left) and 6 social (s1 – s6; right) stimuli during social fear extinction (day 2). **D.** Percentage of investigation time of the 6 social stimuli (s1 – s6) during recall (day 3). **E.** Percentage of vigilance time during social fear extinction of the 3 non-social (ns1, ns3; left) and 6 social stimuli (s1 – s6; right). **F.** Percentage of vigilance time of the 6 social stimuli (s1 – s6) during recall (day 3). Data represent means ± SEM. ## p < 0.01 SFC^+^/males vs SFC^−^/males (C, E), $ p ≤ 0.05 SFC^+^/females vs SFC^−^/females (C, E), ** p < 0.01 SFC^+^/males vs SFC^+^/females (E), (*) p < 0.07 SFC^+^/males vs SFC^−^/females (F), n = 5-7 animals. Analysis was performed using a two-sided unpaired t-test, Mann–Whitney U test (B), or 2-way ANOVA (C-F). Detailed statistics are provided in Supplementary Tables T1 (1-7) and T2 (1-7).

### Astrocytic morphology and calcium levels in the LSc are altered in response to social fear acquisition

The LS is a heterogeneous brain structure with a specific expression pattern for several markers ^16, 53^ and cells dynamically changing their activity during the SFC paradigm ^17^. To assess a potential difference in astrocyte presence between LSr and LSc, we performed a co-immunolabelling of both GFAP and S100β, two well-characterized and widely used astrocytic markers ^54^. We found a higher number GFAP/S100β^+^ astrocytes in the LSc than in the LSr, as well as significant sex differences (Fig. 2A-C). Specifically, female mice exhibited a higher number of astrocytes in comparison to males, irrespective of the specific subsection of LS (LSr: p = 0.006; LSc: p = 0.012; Fig. 2C). Within each sex, an overall higher density of astrocytes was found in the LSc compared to the LSr in male (p = 0.042) and with a tendency in female (p = 0.09) mice (Fig. 2C).

**Figure 2.**
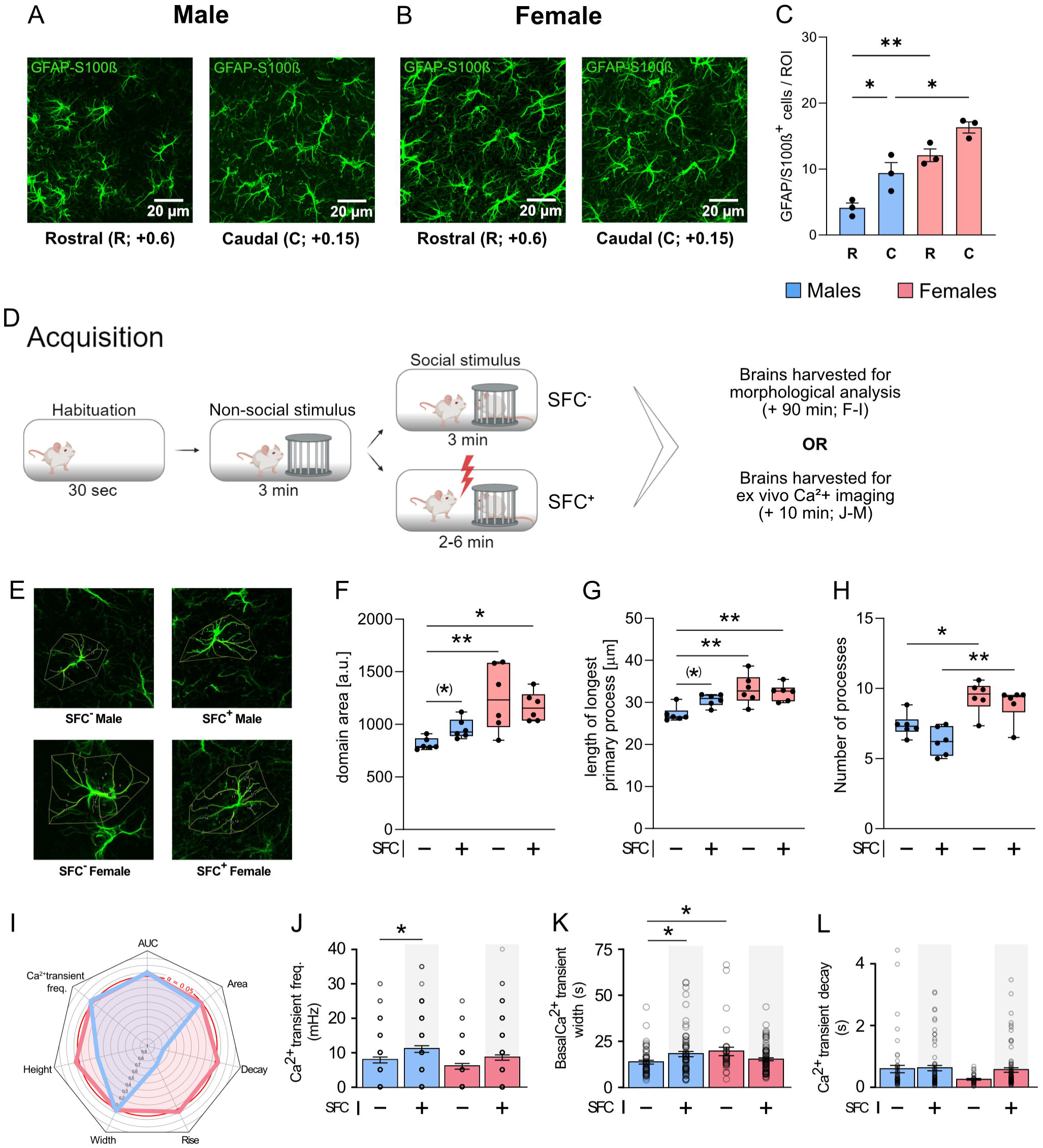
Distribution of GFAP and S100ß expressing (GFAP/S100ß^+^) astrocytes along the rostro-caudal axis of the lateral septum (LS) and the response of LS astrocytes (Morphological alteration: F-I; Calcium response: J-N) to social fear acquisition in male (blue) and female (red) mice. A-B. Representative image from the rostral LS (LSr; Bregma: +0.7) and caudal LS (LSc; Bregma: +0.15) showing immunohistochemistry of GFAP/S100ß^+^ cells (green) of male (A) and female (B) mice. **C.** Number of GFAP/S100ß^+^ cells in the LSr and the LSc of male and female mice. **D.** Scheme of the experimental protocol. **E.** Area of the LSc wherein the morphological and ex vivo calcium (Ca^2+^) imaging was performed. **F.** Quantification of the domain area of LSc astrocytes in male and female mice. **G.** Quantification of the length of the longest primary process in LSc astrocytes of male and female mice. **H.** Quantification of the number of processes in the LSc astrocytes in male and female mice **I.** Spider plot showing the p-value of the comparison between SFC^−^ and SFC^+^ animals for different Ca^2+^ signaling parameters. **J.** Bar plots showing the mean frequency of Ca^2+^ transients triggered by SFC in males and females. **K.** Bar plots showing the mean width of Ca^2+^ transients triggered by SFC in males and females. **L.** Bar plots showing the mean decay time of Ca^2+^ transients triggered by SFC in males and females. Data represent means ± SEM. * p < 0.05 (C, F, H, J, K); ** p < 0.01 (C, F, G, H); (*) p < 0.05 (separate statistics using two-sided unpaired t-test; F, G). Dots represent individual mice (C, F, G, H) and individual cells (J, K, L). Analysis was performed using a two-sided unpaired t-test or a 2-way ANOVA. Detailed statistics are provided in Supplementary Tables T3 (1-5).

Given the high level of morphological plasticity exhibited by astrocytes ^26^, we examined morphological changes of the LSc astrocytes after exposure to the SFC paradigm (Fig. 2D-G). Thus, we performed a GFAP labelling and quantified the GFAP area and the longest GFAP filament in individual astrocytes as previously described ^45^. Interestingly, astrocytes in the LSc of female mice had a higher domain area (Fig. 2F), longer primary processes (Fig. 2G), and a higher number of primary processes (Fig. 2H) in comparison to male mice. However, only in males, separate statistics revealed that social fear acquisition increased the astrocytic domain area (p = 0.011, SFC^+^ vs. SFC^−^; Fig. 2F) and primary process length (p = 0.049, SFC^+^ vs. SFC^−^; Fig. 2G) in the LSc. In contrast, in female SFC^+^ mice, social fear acquisition did not result in comparable changes in astrocytic morphology (Fig. 2F-G), suggesting that the higher number of astrocytes (Fig. 2C) and more complex morphology identified in the LSc of females (Fig. 2F-H) potentially compensates for the lack of discernible morphological responses to social fear acquisition.

Astrocytic Ca^2+^ dynamics are highly heterogeneous in terms of kinetics ^21^. Therefore, we analysed Ca^2+^ dynamics in LSc astrocytes from SFC^−^ and SFC^+^ mice of both sexes under basal conditions (Fig. 2I-L; Fig. S4). In males, the transient calcium responses of astrocytes were more frequent (p = 0.022; Fig. 2J) and larger (p = 0.032; Fig. 2K) in SFC^+^ compared with SFC^−^ mice. However, in SFC^+^ females, the astrocyte calcium transients only tended to be more frequent (p = 0.075; Fig. 2J), shorter (p = 0.074; Fig. 2K), and to have a slower decay (p = 0.061; Fig. 2L). Sex-dependent effects were only found in SFC^−^, but not SFC^+^ mice, with larger and faster calcium transients in females (Fig. 2K-L). Hence, it appears that social fear acquisition influences both the morphology and the calcium transients of LSc astrocytes, depending on the sex of the animals.

### L-AAA-induced inhibition of astrocytic function within the LSc facilitates social fear extinction only in male mice

Given the observed sex differences in social fear extinction (Fig. 1) and in LS astrocytic numbers, morphology, and Ca^2+^ activity (Fig. 2) following social fear acquisition, we aimed to assess the differential role of LS astrocytes in social fear expression in male and female mice. To do so, we first blocked their functions by disrupting the glutamate-glutamine cycle using local L-AAA infusions (Fig. 3). The reported effects of L-AAA include increased glutamate availability, excitotoxicity resulting from NMDA/AMPA receptor overactivation, oxidative stress, and cellular damage ^55, 56^. In a behavioural context, L-AAA has been found to reduce astrocytic support of synaptic plasticity necessary for the consolidation of non-social fear memory, thus resulting in facilitated extinction ^57, 58^. In line with this and other ^43, 44^ studies L-AAA treatment effectively reduced local GFAP signal (p < 0.001 vs vehicle; Fig. S1A, B) without affecting the basal activity of the neuronal network (Fig. S1C). Treatment with L-AAA three days before social fear acquisition did not alter the number of CS-US pairings required by males to acquire social fear (p = 0.16 vs. Veh; Fig. 3C). Interestingly, the same treatment increased the number of CS-US pairings required by the female mice to acquire social fear (p = 0.003 vs. vehicle; Fig. 3F).

**Figure 3.**
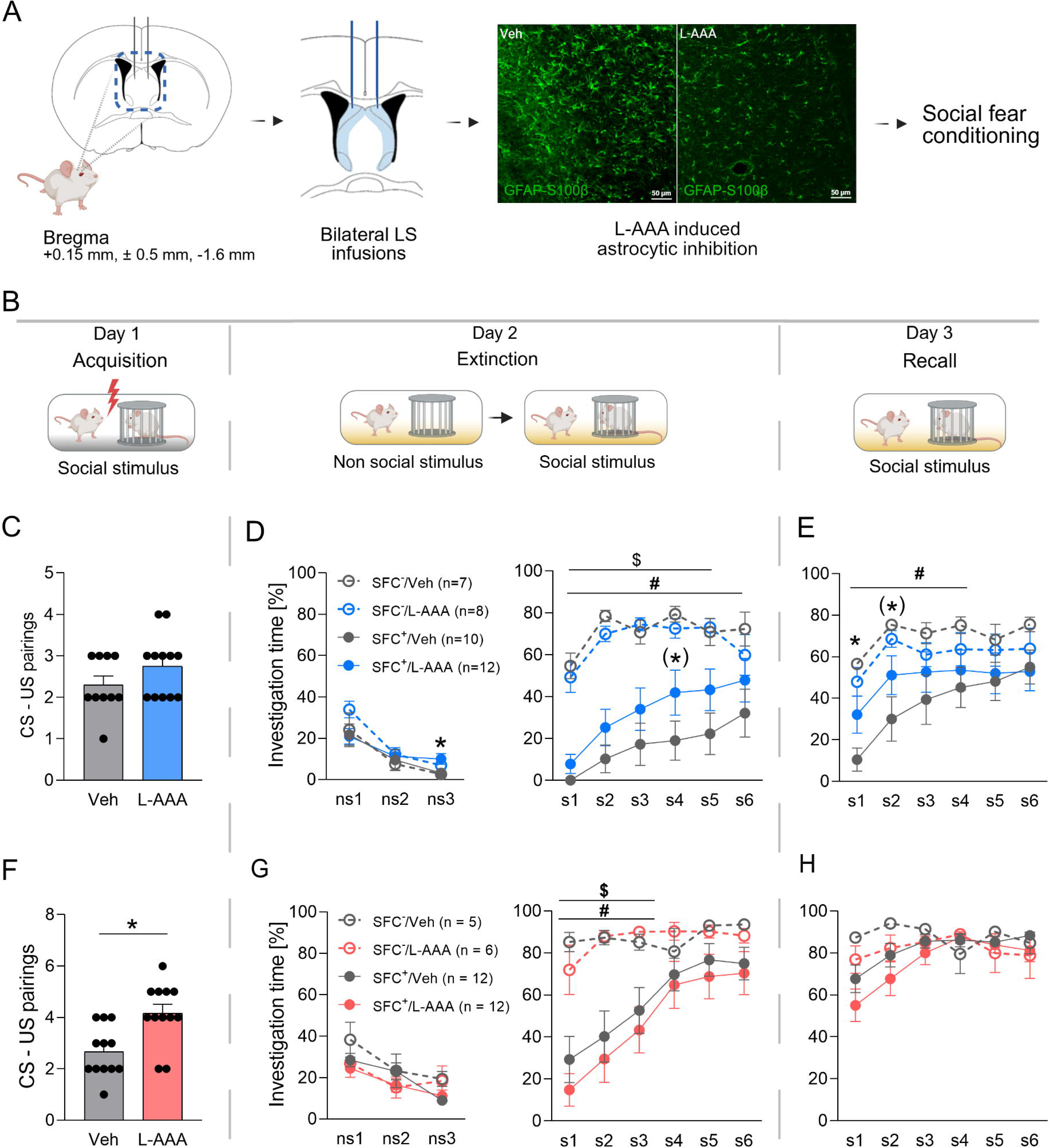
Effects of L-AAA-induced astrocytic inhibition during social fear acquisition (C, F), extinction (D, G), and recall (E, H) in male (blue) and female (red) mice. **A-B.** Schematic presentation of the experimental protocol. **C-F.** Number of CS-US pairings presented to conditioned mice (SFC^+^) during social fear acquisition (day 1) in male and female mice. **D-G.** Percentage of investigation time of the 3 non-social (ns1 - ns3; left) and six social (s1 – s6; right) stimuli during social fear extinction (day 2) in male and female mice. **E-H.** Percentage of investigation time of the six social stimuli (s1 – s6) during recall (day 3) in male and female mice. Data represent means ± SEM. (*) p < 0.07 SFC^+^/L-AAA vs SFC^+^/Veh (D, E), * p ≤ 0.05 SFC^+^/L-AAA vs SFC^+^/Veh (E, F), # p ≤ 0.05 SFC^+^/Veh vs SFC^−^/Veh (D, E, G), $ p ≤ 0.05 SFC^+^/L-AAA vs SFC^−^/L-AAA (D). n = 7-12 animals. Analysis was performed using a two-sided unpaired t-test or a 2-way ANOVA. Detailed statistics are provided in Supplementary Tables T4 (1-7) and T5 (1-7).

**Figure 4.**
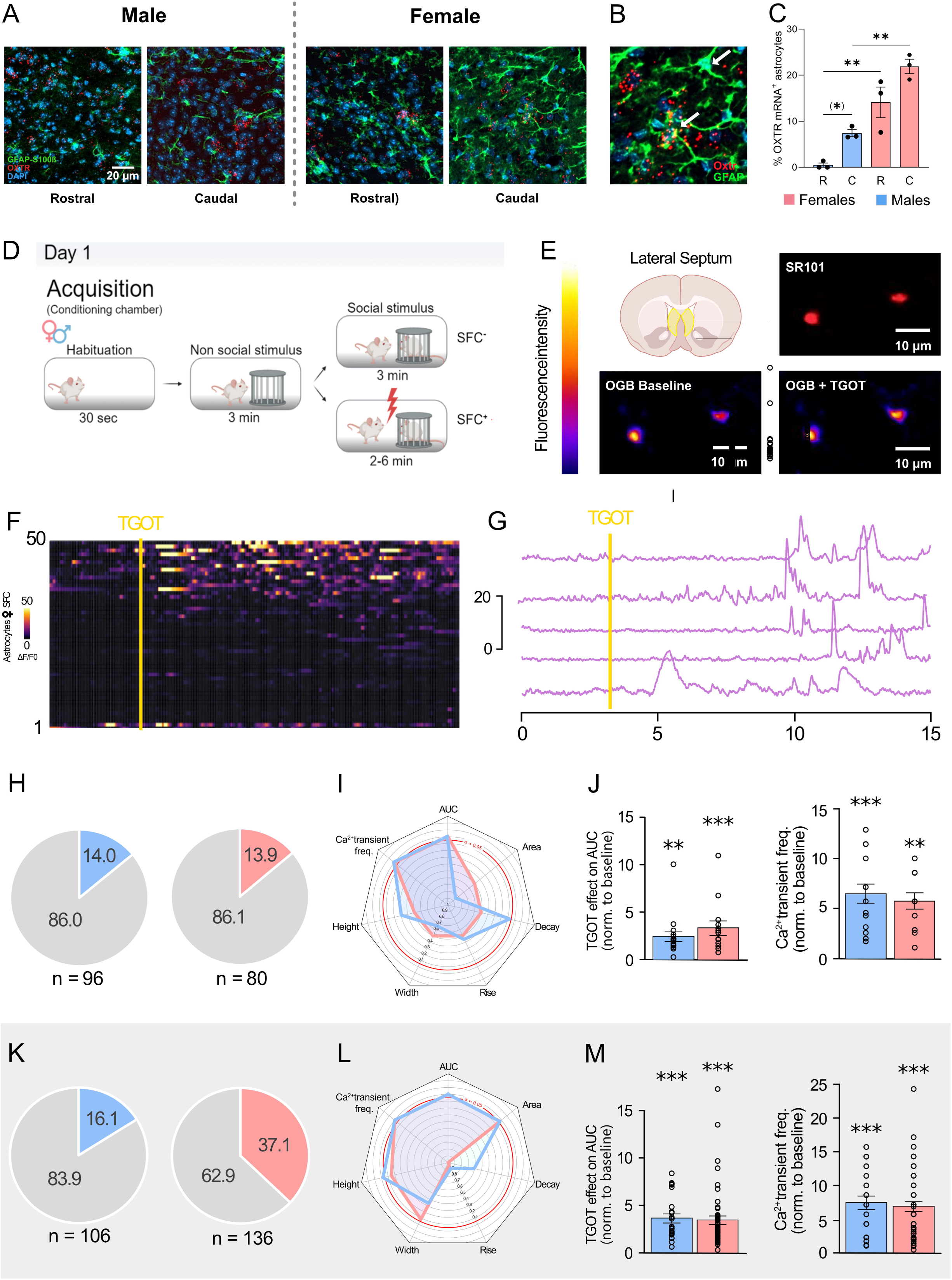
Distribution of oxytocin receptor mRNA-expressing astrocytes (OXTR-GFAP-S100β^+^) along the rostro-caudal axis of the lateral septum (LS) of male (blue) and female (red) mice (A-C) and the response of OXTR-GFAP-S100β^+^ cells in the LS to social fear acquisition and OXTR activation in unconditioned (SFC^−^; H-J) and conditioned (SFC^+^; grey background; K-M) male and female mice. **A.** Representative image from the rostral LS (R; Bregma: +0.7) and caudal LS (C; Bregma: +0.15) showing OXTR-GFAP/S100ß^+^ cells. **B.** High-magnification image showing astrocytes with and without for OXTR. **C.** Comparison of OXTR-GFAP/S100ß^+^ cells in LS. **D**. Scheme of experimental setups used for assessment of SFC. **E**. Images of LS astrocytes identified through SR101 (red, top right) and corresponding pseudo-color images of OGB1 fluorescence during baseline and after drug application (bottom left and right, stacks of 20 images over 10 seconds of recording. Scale bar, 10 µm. **F**. Heatmap of OGB1 fluorescence intensity of individual astrocytes in response to TGOT (500nM). The vertical line indicates the 20s of TGOT application. **G**. Example traces of OGB1 fluorescence intensity of individual astrocytes in response to TGOT (500nM). The vertical line indicates the 20s of TGOT application. **H-J**. Astrocytes calcium transients in SFC- mice. **H**. Proportion of LS astrocytes responding to TGOT (500nM) application in SFC^−^ male and female mice (n_cell male_=96, n_record male_=33, n_mouse male_=3; n_cell female_=80, n_record female_ = 29, n_mouse female_=3). **I.** Comparison of calcium activity triggered by TGOT (500nM) application in SFC^−^ male and female mice. Spider plots showing the p-value of the comparison between the basal state and after the application of TGOT for different calcium signaling parameters. **J**. Bar plots showing mean AUC and mean frequency of calcium transients triggered by TGOT application (500µM) in male and female SFC^−^ mice normalized to baseline values (n_cell male_=96, n_record male_=33, n_mouse male_=3; n_cell female_=80, n_record female_=29, n_mouse female_=3). **K-M**. Astrocytes calcium transients in SFC+ mice. **K**. Proportion of LS astrocytes responding to TGOT (500nM) application in male and female SFC^+^ mice (n_cell male_=106, n_record male_=33, n_mouse male_=4; n_cell female_=136, n_record female_=46, n_mouse female_=4). **L**. Comparison of calcium activity triggered by TGOT (500nM) application in male and female SFC^+^ mice. Spider plots showing the p-value of the comparison between the basal state and after the application of TGOT for different calcium signaling parameters. **M**. Bar plots showing mean AUC and mean frequency of calcium transients triggered by TGOT application (500nM) in male and female SFC^+^ mice normalized to baseline values (n_cell male_=106, n_record male_=33, n_mouse male_=4, n_cell female_=136, n_record female_= 46, n_mouse female_=4). Data are expressed as mean across animal ± SEM. * p<0.05, ** p<0.01, *** p<0,001, two-sided unpaired t-test, Mann-Whitney U test, or Anova 2. Detailed statistics are provided in Supplementary Tables T6 and T7.

During social fear extinction on day 2 (Fig. 3D, G), male, but not female, SFC^+^/L-AAA mice showed higher investigation (p = 0.035) and lower vigilance (p = 0.031) times of the 3^rd^ non-social stimulus compared to SFC^+^/Veh mice (Fig. 3D, G and S3C, E left), indicating a mild reduction in general anxiety. During exposure to s1, all male and female SFC^+^ mice expressed social fear, independent of treatment, as indicated by lower social investigation and higher vigilance behavior compared to their respective SFC^−^ mice (Fig. 3D, G, and S3C, E right). The expression of social fear gradually decreased over the extinction period in all groups. Interestingly, astrocytic inhibition in the LSc accelerated social fear extinction in males, but not females, as indicated by higher investigation times at s4 (p = 0.061 SFC^+^/L-AAA vs. SFC^+^/Veh mice; Fig. 3D right). In support, while male SFC^+^/Veh mice still showed lower investigation of s6 (p = 0.016 vs SFC^−^/Veh group), male SFC^+^/L-AAA mice did not differ from SFC^−^ mice at this time point (p = 0.418 vs SFC^−^/L-AAA). In females, L-AAA had no effect on social exploration.

During recall on day 3 (Fig. 3E, H), the facilitatory effect of L-AAA on extinction in male mice was still evident, as SFC^+^/L-AAA males displayed higher social investigation times (s1-s4, p < 0.05; Fig. 3E) and lower vigilance times (s1-s4, p < 0.05; Fig. S3D) compared to SFC^+^/Veh mice. However, in female mice, no differences in social investigation (Fig. 3H) or social vigilance (Fig. S3F) were observed during recall, suggesting that an overall inhibition of astrocytic function does not alter social fear extinction in female mice.

### OXTR^+^ astrocytes show a sex-dependent distribution pattern in the LS and exhibit altered Ca^2+^ response to OXTR agonism after social fear acquisition

We have previously demonstrated that OXT signalling in the LS plays a critical role in facilitating social fear extinction in both male ^13, 59^ and female ^9^ mice. To reveal the specific contribution of astrocytic OXTR, we first analyzed the distribution of astrocytic OXTR within the LS using in-situ hybridization of the OXTR mRNA, together with a co-immunostaining for GFAP and S100B (Figure 4A-B). Analysis of the percentage of OXTR^+^ astrocytes within the rostral or caudal part of the LS revealed sex- (F_1,8_ = 55.19, p < 0.001) and region- (F_1,8_ = 15.32, p = 0.003) specific distribution patterns. In detail, females showed a higher percentage of OXTR^+^ astrocytes in both the LSr (p = 0.004) and the LSc (p = 0.003) compared to males (Fig. 4A-C). In terms of anatomical distribution, a higher percentage of OXTR^+^ astrocytes was found in the LSc compared to the LSr (p = 0.003; Fig. 4C) in male mice while no such differences could be observed in females.

Given the significant differences observed between male and female OXTR astrocytes, we analysed the responses of astrocytes to OXTR activation using slice calcium imaging in SFC^−^and SFC^+^ male and female animals (Fig. 4D-M; Fig. S5). In SFC^−^ mice of both sexes (Fig. 4H-J), a similar proportion of astrocytes responded to TGOT application (p = 1.0; Fig. 4H). This lack of sex differences in astrocytic responses was consistent across all analysed parameters, including AUC and calcium transient frequencies (Fig. 4I-J; Fig S5). In contrast, in SFC^+^ mice (Fig. 4K-M; Fig. S5), a larger proportion of astrocytes responded to the application of TGOT in females than in males (p = 0.0128; Fig. 4K). However, the kinetics of response were similar across all analysed parameters, including AUC and calcium transient frequency, between male and female SFC^+^ (Fig. 4L-M; Fig. S5). Of importance, in males, SFC did not modify the calcium response of astrocytes, neither in the proportion of astrocytes responding to TGOT in males (p = 0.986; Fig. 4H and 4K), nor in the calcium transient parameters (Fig. 4J and 4M; Fig. S5). However, in females, SFC induced a TGOT response in an increased proportion of astrocytes (p = 0.012; Fig. 4H and 4K), but no differences were detected in the calcium transient parameters (Fig. 4J and 4M; Fig. S5). In conclusion, OXTR activation leads to a similar astrocyte response in male and female SFC^−^mice, but not in SFC^+^ mice, in which female mice appear more sensitive to OXTR agonism.

### Knockdown of astrocytic OXTR in the LS facilitates the extinction of social fear in male and female mice

Based on the finding of an elevated astrocytic calcium response to TGOT in the LS of SFC^+^ mice of both sexes (Fig. 4), we investigated whether astrocytic OXT signaling within the LSc is involved in the behavioral responses during social fear acquisition, extinction, and recall. For astrocytic-specific functional manipulation, we used a viral construct expressing a shRNA to exclusively knock down astrocytic OXTR (OXTR-shRNA) ^29^. OXTR-shRNA significantly downregulated relative OXTR mRNA in LSc punches of females (p = 0.023 vs. scr; Fig. S6C), but not of males (p = 0.074 vs. scr; Fig. S6B), probably due to the extremely low baseline OXTR mRNA expression identified in males (Fig. 4A-B).

In both male and female mice, functional knockdown of OXTR did neither affect social fear acquisition (day 1, Fig. 5C and F) nor the investigation of the three non-social stimuli (day 2, Fig. 5D and S6E left; 5G and S6G left). During social fear extinction training, all male and female SFC^+^ mice expressed social fear independent of treatment, indicated by significantly lower social investigation and higher vigilance times during exposure to s1 compared to respective SFC^−^ controls (Fig. 5D and S6E right; 5G and S6G right). No overall treatment effect could be observed on social investigation. However, separate statistics for each time point of extinction revealed that OXTR knockdown accelerated social fear extinction in both male and female SFC^+^ mice, as indicated by higher investigation times at s5 (males: p = 0.028, females: p = 0.03 vs. SFC^+^/scr; Fig. 5D and Fig. 5G right). In support of a faster extinction after astrocytic OXTR knockdown in males, SFC^+^/scr mice still showed lower investigation of s5 and s6 (s5 p = 0.004; s6 p = 0.053, vs. SFC^−^/scr mice; Fig. 5D right), while male SFC^+^/OXTR^−^ did not differ at these time points (p > 0.05 vs SFC^−^/OXTR^−^; Fig. 5D right) from their respective SFC^−^ controls. Similarly, female SFC^+^/scr mice still showed lower investigation of s6 (p = 0.04 vs. SFC^−^/scr mice; Fig. 5G right), whereas female SFC^+^/OXTR^−^did not differ at this time point (p = 0.59 vs. SFC^−^/scr mice; Fig. 5G right). In line with the above data, we found that OXTR knockdown reduced vigilance times in SFC^+^ male mice during s4 to s6 (p > 0.05 vs. SFC^−^/scr; Fig. S6E) and in SFC^+^ female mice during exposure to s1, s4, and s5 (s1 p = 0.02; s4 p = 0.012; s5 p = 0.016 vs. SFC^+^/scr mice; Fig. S6G).

**Figure 5.**
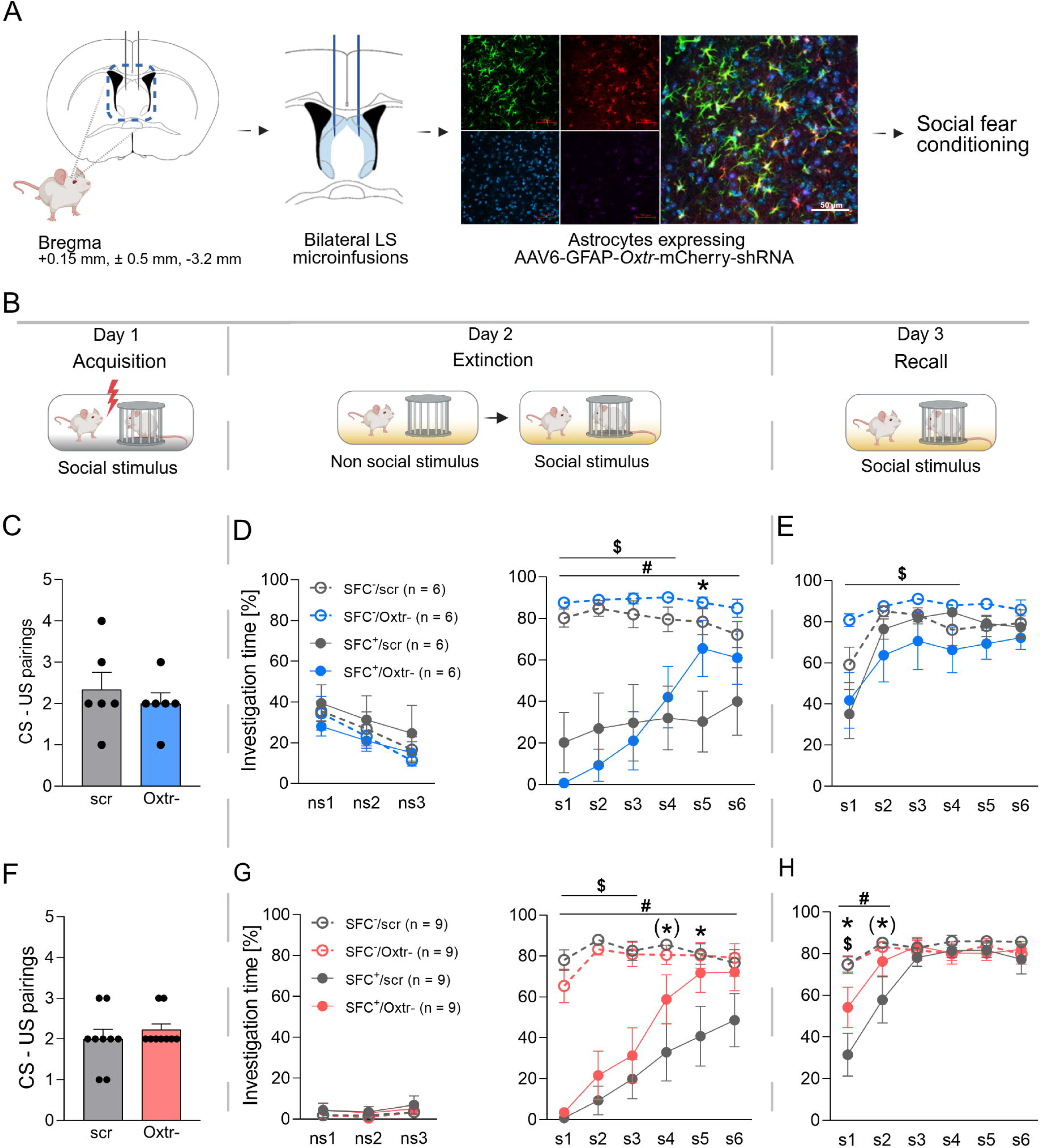
Effects of OXTR mRNA knockdown (OXTR^−^) in caudal lateral septum (LSc) astrocytes on social fear acquisition (C, F), extinction (D, G), and recall (E, H) in male (blue) and female (red) mice. **A-B.** Schematic presentation of the experimental protocol. The LSc of male and female mice were infused with an AAV expressing an shRNA targeting OXTR mRNA (OXTR^−^) or an AAV expressing a scrambled control (scr) three weeks prior to the SFC paradigm. **C-F.** Number of CS-US pairings presented to conditioned mice (SFC^+^) during social fear acquisition (day 1) in male and female mice. **D-G.** Percentage of investigation time of the three non-social (ns1 - ns3; left) and six social (s1 – s6; right) stimuli during social fear extinction (day 2) in male and female mice. **E-H.** Percentage of investigation time of the six social stimuli (s1 – s6) during recall (day 3) in male and female mice. Data represent means ± SEM. * p ≤ 0.05 SFC^+^/OXTR^−^ vs SFC^+^/Scr (D, G, H), # p ≤ 0.05 SFC^+^/Scr vs SFC^−^/Scr (D, G, H), $ p ≤ 0.05 SFC^+^/OXTR^−^ vs SFC^−^/OXTR^−-^ (D, G, H), (*) p < 0.07 SFC^+^/OXTR^−^ vs SFC^+^/Scr (G, H). Detailed statistics are provided in Supplementary Tables T8 (1-7) and T9 (1-7).

During recall on day 3, the effects of OXTR^−^ seen in females during extinction were still evident with higher social investigation times (s1 p = 0.042; s2 p = 0.060; Fig. 5H) and lower vigilance times (s1 p = 0.005; s2 p = 0.034; Fig. S6H) compared to SFC^+^/scr. Interestingly, in male mice, while no treatment effect was found in the investigation or vigilance times of SFC^+^ mice, SFC^+^/OXTR^−^ showed lower investigation times of s1 to s4 (s1 p = 0.011; s2 p = 0.032; s3 p = 0.068; s4 p = 0.040; Fig. 5E) and higher vigilance times of s1 (p = 0.009; Fig. S6F) compared to SFC^−^/OXTR^−^ mice. Overall, these findings suggest that astrocytic OXTR mRNA knockdown in the LSc of both male and female mice does not affect the acquisition of social fear, but facilitates its extinction.

## Discussion

In the present study, we analysed the sex-dependent involvement of astrocytes within the LS, particularly of OXTR^+^ astrocytes, in the acquisition and extinction of social fear using the SFC paradigm. A direct behavioural comparison of male and female SFC^+^ mice revealed elevated social fear expression and slower social fear extinction in males. Also, male mice have a generally lower astrocyte density within the LS, including the LSc, where astrocyte abundance is highest compared to the LSr. Only in males, social fear acquisition resulted in morphological changes in LSc astrocytes, whereas an enhanced baseline astrocytic Ca^2+^ activity was found in both male and female SFC^+^ mice after acquisition. The indication for a sex-specific functional involvement of LS astrocytes in social fear extinction comes from pharmacological inhibition of astrocytic functions, which facilitated extinction only in male SFC^+^ mice. With respect to astrocytic OXT signaling, we identified a lower number of OXTR^+^ astrocytes in the LS of males, which was, nevertheless, highest in the LSc compared to the LSr in both sexes. Ex vivo Ca^2+^ imaging showed that social fear acquisition enhanced the response of LSc astrocytes to OXTR agonism in both sexes. Further, astrocyte-specific knockdown of OXTR in the LSc facilitated the extinction of social fear in both male and female mice. Thus, OXTR^+^ astrocytes in the mouse LSc are an important component of circuits underlying the regulation of social fear and rather contribute to sustained social fear after social trauma.

### Striking sex differences in social fear expression, and in astrocytic distribution in the LS

A direct comparison of male and female mice in the SFC paradigm revealed subtle differences during extinction, with slower social fear extinction in male SFC^+^ mice (Fig. 1C, 1E). Sex differences in behavioral responses have been repeatedly reported in rodents in the context of fear and anxiety ^60–62^, likely based on differences in sex steroid levels ^63^ and of various social behaviors ^53, 64, 65^. Thus, males may perceive the social stimuli exposed to during fear extinction as a threat due to their higher territorial aggression level ^66^, whereas female rodents display stronger social preference behavior and higher social motivation ^67–69^.

We focused on astrocytic functions in the LS, a region important for the regulation of socio-emotional behaviours ^15, 16^, specifically for modulating the extinction of social fear ^9, 13, 14, 17, 19, 59^. We identified a distinct astrocyte distribution pattern, with the highest astrocytic density in the LSc compared with the LSr of both sexes (Fig. 2C). As shown in selected brain regions, a higher density of astrocytes has been linked with supporting heightened neuronal demand ^54, 70^, while chemogenetic silencing of astrocytes hinders neuronal synchronisation ^71^. Thus, the relatively high astrocytic density in the LSc in both males and females suggests their local importance in regulating and coordinating neuronal responses and, subsequently, socio-emotional behavior. We could also confirm ^24^ a higher astrocytic density in the LSc of females (Fig. 2C), accompanied by a larger astrocytic domain area (Fig. 2F), longer primary processes (Fig. 2G), and a larger number of processes (Fig. 2H). Sexual dimorphism in astrocytic density was found to be region-specific in the mouse brain, with males exhibiting a higher density than females in the medial amygdala ^72^, but not in the hippocampus ^40^. Such region-dependent sexual dimorphism in astrocytic density and activity has been linked to differential processing of fear memory ^40^.

### Astrocytic plasticity upon social fear acquisition

Interestingly, in response to acquisition, LSc astrocytes showed sex-dependent morphological changes, with an enhanced GFAP-positive astrocytic domain area observed only in male (Fig. 2F and 2G), but not female (Fig. 2F and 2G) SFC^+^ mice. This result is in agreement with a sex-dependent astrocytic remodeling seen after exposure to an acute stressor and suggests an overall decrease in the morphological complexity of astrocytes ^26^. Given that perisynaptic astrocytic processes are responsible for neurotransmitter uptake and ionic homeostasis control ^57, 73^, it is likely that the observed GFAP structural change reflects a modulation of the synaptic strength ^26^ and/or a synaptic remodeling that facilitates learning during social fear acquisition. However, to what extent this morphological adaptation of astrocytes in males compensates for their lower astrocytic number and density identified in the LSc (Fig. 2C) remains to be shown. Sex-dependent differences in astrocytic morphological responses may contribute to the observed behavioural differences in social fear processing. Similar sex-dependent dimorphic astrocytic alterations have been shown across multiple brain regions in response to chronic stress ^50^ and within the hippocampus in the context of spatial memory ^40^.

Social fear acquisition not only affected social behaviour and astrocytic morphology, but also ex vivo astrocytic Ca²⁺ activity in a sex-dependent way. Following social fear acquisition, we identified an elevated Ca²⁺ activity of LS astrocytes of SFC^+^ compared to SFC^−^ male mice (Fig. 2J-2K), an effect that was absent in females. The heightened Ca^2+^ activity was reflected by an increased frequency and amplitude of Ca²⁺ transients. While this might reflect a functional adaptation to the observed morphological change (Fig 2A-H), it remains to be shown whether this indicates a compensatory mechanism in males with lower astrocyte density in the LS. In addition, such a change in calcium activity might reflect a greater intrinsic astrocytic modulatory role at surrounding synapses, potentially leading to overall modulation of neuronal excitability and enhanced network synchronization ^74, 75^. Thus, an increase in astrocytic domain area in response to social fear acquisition in males may facilitate synaptic plasticity in the LSc, thereby enhancing consolidation of social fear memory and accounting for the observed impairment of extinction in males. In contrast, in females, the generally higher astrocytic density in the LSc might sufficiently mediate the neuronal plasticity and consequent consolidation of social fear memory post-acquisition.

### Involvement of LS astrocytes in social fear expression and extinction

Further emphasizing the differential role of LS astrocytes between female and male behavioural responses, pharmacological inhibition of local astrocytic activity by disrupting the glutamate-glutamine cycle with L-AAA significantly reduced social fear expression and facilitated social fear extinction, but only in male mice (Figs. 3D, S3C). The behavioural effect of L-AAA was highly context-specific, as it did not alter naturally occurring social preference (Fig S2D), social novelty preference (Fig. S2E), or anxiety-related behaviours (Fig. S2A-C). In contrast to males, L-AAA-induced astrocytic inhibition did not affect social fear expression and extinction in female mice (Fig. 3G), but rather delayed social fear acquisition (Fig 3F). These differential results in males and females generally show that astrocytic actions within the LS make animals more susceptible to social trauma (in females) and more resistant against extinction training-induced reversal of social fear (in males). However, the mechanisms underlying the sex-specific behavioural effects of astrocytic inhibition on social fear acquisition and extinction in females and males, respectively, including the contribution of sex hormones, particularly estrogen ^76–80^ remain to be studied.

### Facilitation of social fear extinction by LS astrocytic OXTR

OXT signaling within the LS is essential for social fear extinction in both male and female mice ^9, 13^. Here, we identified OXTR^+^ astrocytes in LS, with a distribution pattern mirroring overall higher astrocyte density in the LSc than the LSr (Fig. 4C), and higher astrocyte density in females compared to males (Fig. 4C). In line, sex differences in brain OXTR with higher expression and binding in females ^24, 81^, and the promoting effect of estrogens ^10, 36^, are well documented. Regarding the functionality of astrocytic OXTR, treatment with the selective OXTR agonist TGOT elevated astrocytic Ca²⁺ activity in the LSc of both male and female mice (Fig. 4J and 4M), which supports earlier findings ^25, 82, 83^. Interestingly, despite generally lower OXTR expression, SFC^−^ control males displayed similar astrocytic Ca²⁺ activity in response to TGOT compared to SFC^−^ females (Fig. 4H and 4J). However, after social fear acquisition, a higher percentage of astrocytes responded to TGOT in female SFC^+^ animals (Fig. 4K), suggesting that acquisition of social fear led to a sensitization of OXTR^+^ astrocytes in females.

To assess the involvement of LS astrocytic OXTR in social fear acquisition and extinction, we used an OXTR-shRNA exclusively expressed under the control of GFAP ^29^. Bilateral OXTR-shRNA transfection in the LS resulted in a significant cell type-specific OXTR knockdown (approximately 50%) in LS astrocytes, but only in females, (Fig. S6C). This is likely due to the low basal levels astrocytic OXTR expression we identified in the male LS (Fig.4C). However, without altering social fear acquisition, OXTR-shRNA treatment facilitated fear extinction in both male and female mice, but with a more robust effect in females, reflected by increased investigation of the social stimuli and decreased vigilance behavior in SFC^+^/OXTR^−^ mice (Fig. 5D and 5G). Notably, the facilitatory effect of astrocytic OXTR knockdown on social fear extinction was also reflected by higher social investigation times in SFC^+^/OXTR^−^ mice during recall in comparison to SFC^+^/scr controls (Fig. 5E and 5H).

The higher astrocytic OXTR expression and Ca^2+^ response in female LSc may explain why local astrocytic OXTR knockdown has a greater behavioural impact in female mice. Reduced astrocytic OXTR signaling appeared to facilitate the extinction of social fear by diminishing astrocytic actions on synaptic plasticity and transmission, and memory consolidation ^57^ in fear-related circuits, thereby promoting the extinction of social fear. However, these findings challenge the well-established view of OXT as a fear-reducing and prosocial neuropeptide acting within the LS during social fear extinction ^9, 13^. While these effects are likely based on neuronal OXTR-mediated signaling, astrocytic OXTR signaling also contributes to its anxiolytic effects, for example, within the PVN ^26, 29^. Clearly, cell type-specific effects of OXT need to be consistently considered in future studies.

In conclusion, we describe a sex-dependent distribution of astrocytes, specifically of OXTR^+^ astrocytes, in the LS with the highest densities in the LSc, and their sex-dependent involvement in social fear extinction. Inhibition of local astrocytic functions facilitated extinction in male mice, while cell type-specific reduction of LS astrocytic OXTR signaling facilitated extinction in both males and females. Thus, astrocytic and neuronal OXTR-mediated signaling within the LS seem to exert opposite effects in the context of social fear. Future research should explore the underlying molecular and subcellular effects of astrocytic modulation of synaptic plasticity and behavior in response to OXT within social fear-related circuits including the LS.

## Acknowledgements

This work was supported by the German Research Foundation (DFG) grant (Ne465/31; to IDN), the DFG Graduate School (GRK-2174; to IDN, RM), the Centre National de la Recherche Scientifique contract UPR3212, the Université de Strasbourg contract UPR3212, the Agence Nationale de la Recherche (ANR, French Research Foundation) grants (ANR-24-NEU2-0008, ANR-24-CE37-5566, 23-CE37-0015-01, the FRM Equipe grant EQU202403018071; to AC), the Graduate School of Pain EURIDOL (ANR-17-EURE-0022; to AC and CD), and the CNRS International Research Project grant (ICOT2023; to AC)

## Author contribution statements

Project conception, LB, RM, IDN; Methodology, LB, RM, CD, K-YW, HP, BDB, AC, IDN; anatomic analysis: LB, RM; Ex vivo patch-clamp electrophysiology, AC, HP; Ex vivo calcium imaging, CD, K-YW; Behavior analysis: LB, RM; Writing, RM, AC, IDN; Illustrations, RM, CD, AC; Project administration and supervision, AC, IDN.

