## Supplementary Figures for "Sex-dependent involvement of lateral septum astrocytes in social fear: Role of oxytocin receptor signaling"

**S1: Effect of L-AAA infusion on astrocytes in the in caudal lateral septum.**

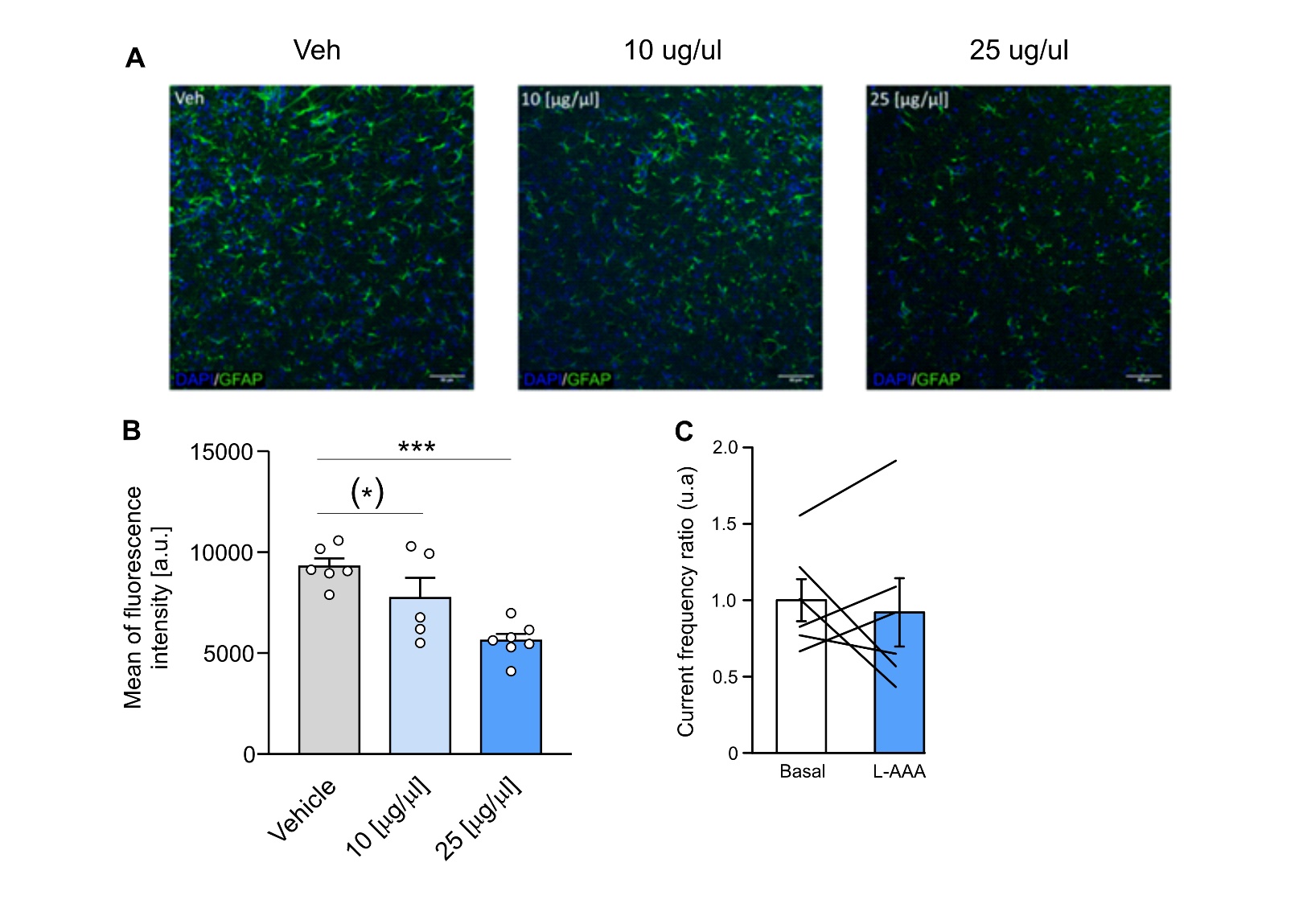

**Figure S1. Effects of L-AAA infusion on astrocytes in the caudal lateral septum (LSc) of male mice. A.** Representative image from the LSc showing immunohistochemical staining for glial fibrillary acidic protein expressing (GFAP, green) and nuclei stained with DAPI (blue) from Vehicle-treated or L-AAA treated (10 µg/µl or 25 µg/µl) mice. Scale bar = 20 µm. **B.** Quantification of mean fluorescence of GFAP in Vehicle, 10 µg/µl, and 25 µg/µl L-AAA treated group. **C.** Ex vivo patch-clamp recordings of spontaneous inhibitory post-synaptic currents (IPSC) recorded in random neurons before and after incubation of the slice in 1mM L-AAA for 45 min. Data represent means ± SEM. (*) p ≤ 0.07, *** p < 0.001. Detailed statistics are provided in Supplementary Tables T10.

**S2: L-AAA-induced inhibition of astrocytic function does not alter anxiety-like behavior, social preference, or social novelty.**

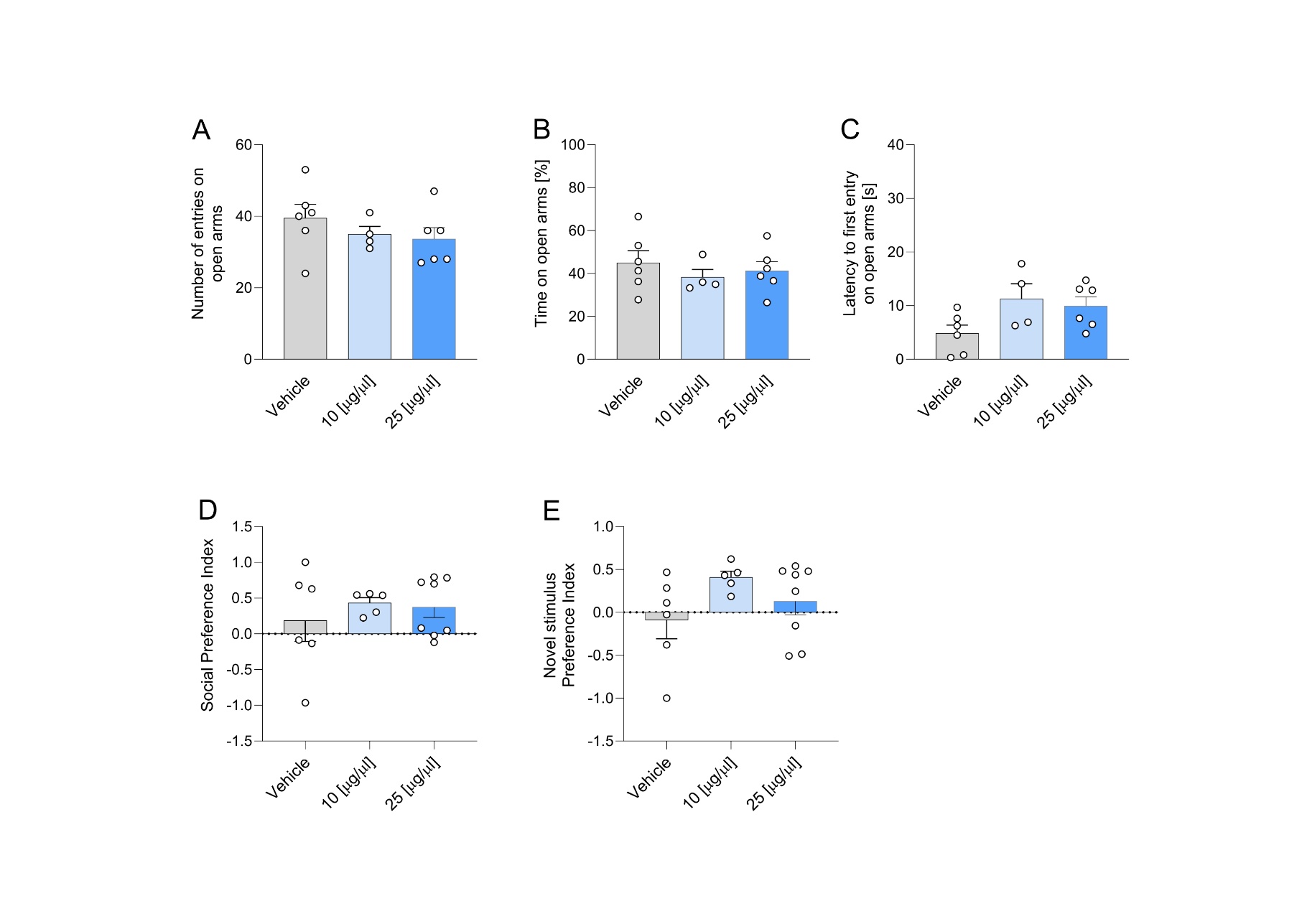

**Figure S2. Effects of L-AAA-induced astrocytic dysfunction in the caudal lateral septum (LSc) in male mice on anxiety-like behavior (A-C), social preference (D), and social novelty (E). A-C.** show quantification of the number of entries (A), the percentage of time spent in open arms (B), and the latency to the first entry into open arms (C), in the Elevated Plus Maze in Veh, 10 µg/µl and 25 µg/µl L-AAA-treated groups. **D-E.** Social Preference Index (D) and Novel stimulus Preference Index (E) in Veh-, 10 µg/µl and 25 µg/µl L-AAA-treated mice. Data represent means ± SEM. Detailed statistics are provided in Supplementary Tables T11.

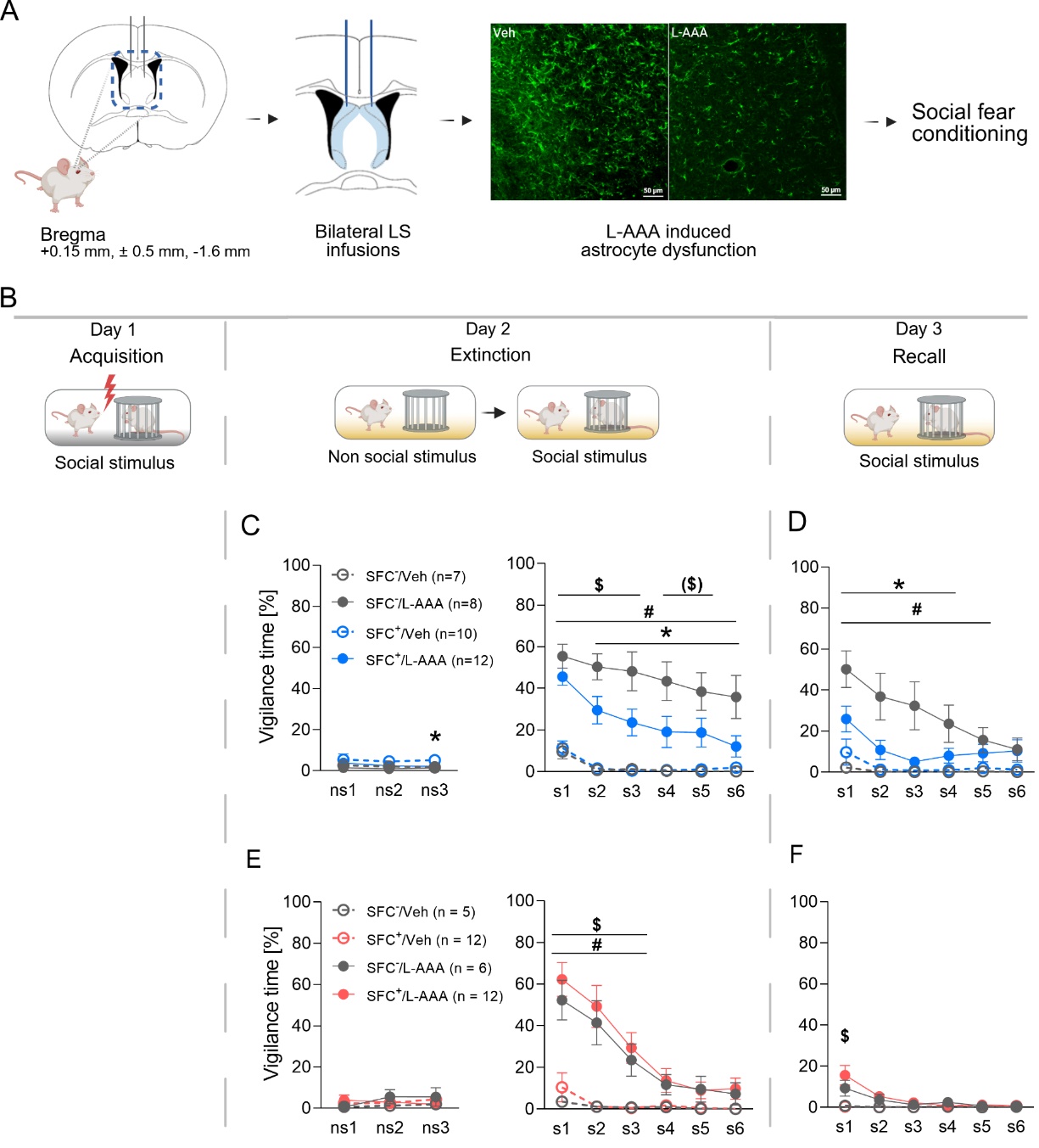
**S3: L-AAA-induced inhibition of astrocytic function within the caudal lateral septum inhibits vigilance during social fear extinction in male mice.**

**Figure S3. Effects of L-AAA-induced astrocytic inhibition on vigilance during social fear extinction (C, E), and recall (D, F) in male (blue) and female (red) mice.** **A-B.** Schematic representation of the surgical and experimental protocol. **C, E.** Percentage of investigation time of the 3 non-social (ns1 - ns3; left) and six social (s1 – s6; right) stimuli during social fear extinction (day 2) in male and female mice. **D, F.** Percentage of investigation time of the six social stimuli (s1 – s6) during recall (day 3) in male and female mice. Data represent means ± SEM. * p ≤ 0.05 SFC^+^/L-AAA vs SFC^+^/Veh (C, D), # p ≤ 0.05 SFC^+^/Veh vs SFC^-^/Veh (C, D, E), $ p ≤ 0.05 SFC^+^/L-AAA vs SFC^-^/L-AAA (C, D, E, F), ($) p ≤ 0.07 SFC^+^/L-AAA vs SFC^-^/L-AAA (C). n = 7-12 animals. Analysis was performed using a two-sided unpaired t-test or a 2-way ANOVA. Detailed statistics are provided in Supplementary Tables T12 (1-7) and T13 (1-7).

**S4: Effect of SFC on basal astrocytic activity in male and female mice.**

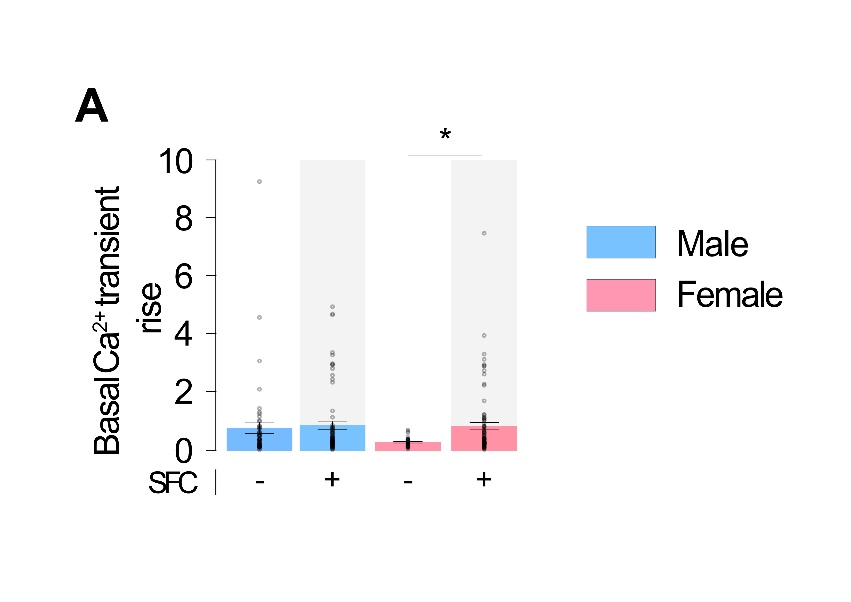

**Figure S4**. A. Astrocytic basal activity in male and female SFC^-^ and SFC^+^ (Grey background) mice. Dots represent individual cells. Data represent means ± SEM. * p ≤ 0.05. Detailed statistics are provided in Supplementary Table T7.

**S5: Effect of TGOT on astrocytic activity of male and female SFC^-^ and SFC^+^ mice.**

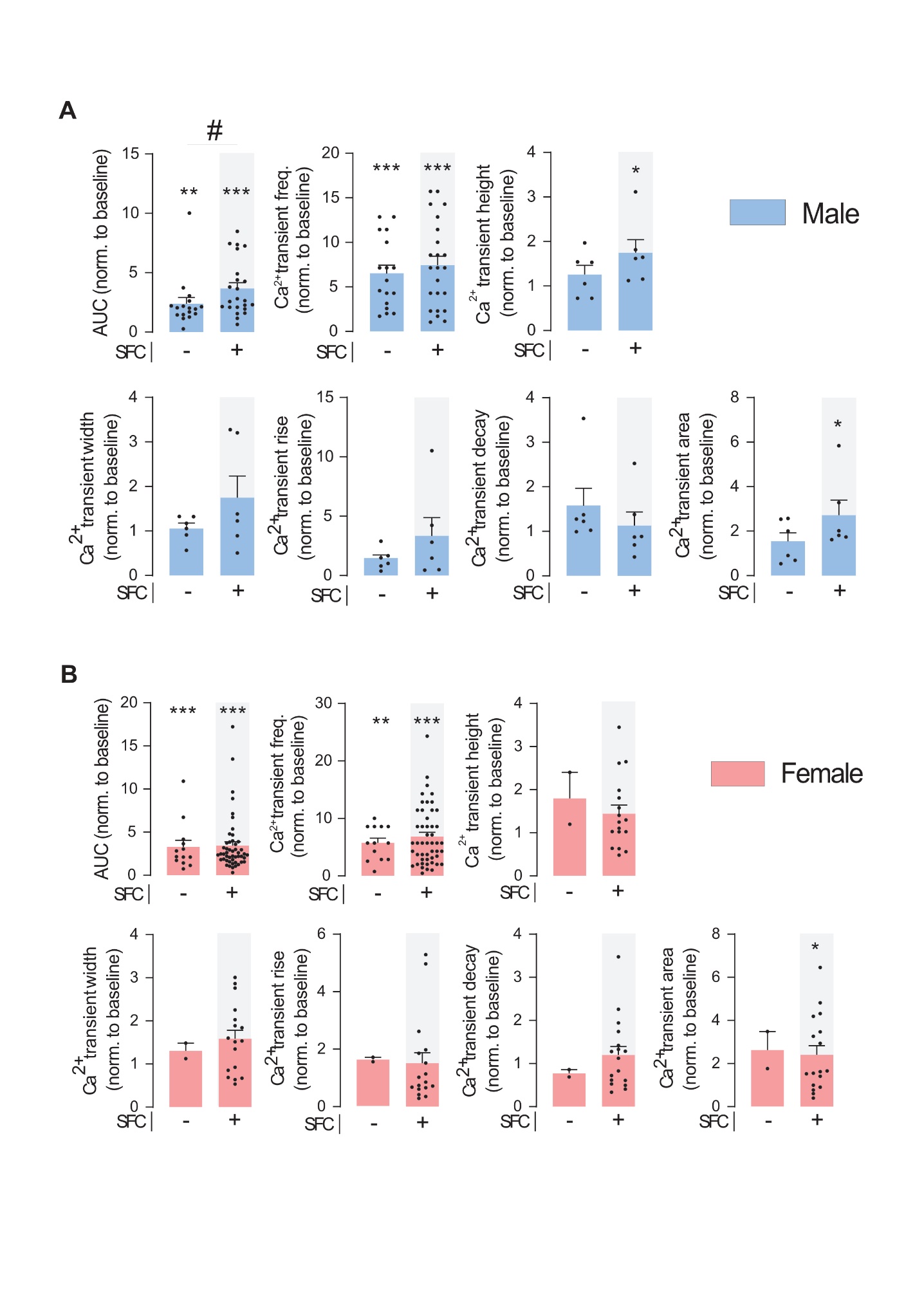

**Figure S5.** **Effect of TGOT on astrocytic activity of male and female SFC^-^ and SFC^+^ (grey background) mice. A.** Astrocytes calcium transients’ characteristics in male mice. **B** Astrocytes calcium transients’ characteristics in female mice**.** Dots represent individual cells. * p ≤ 0.05, ** p ≤ 0.01, *** p ≤ 0.001. Detailed statistics are provided in Supplementary Table T7.

**S6: Knockdown of astrocytic OXTRs in the LS reduces vigilance behavior during the extinction of social fear in male and female mice.**

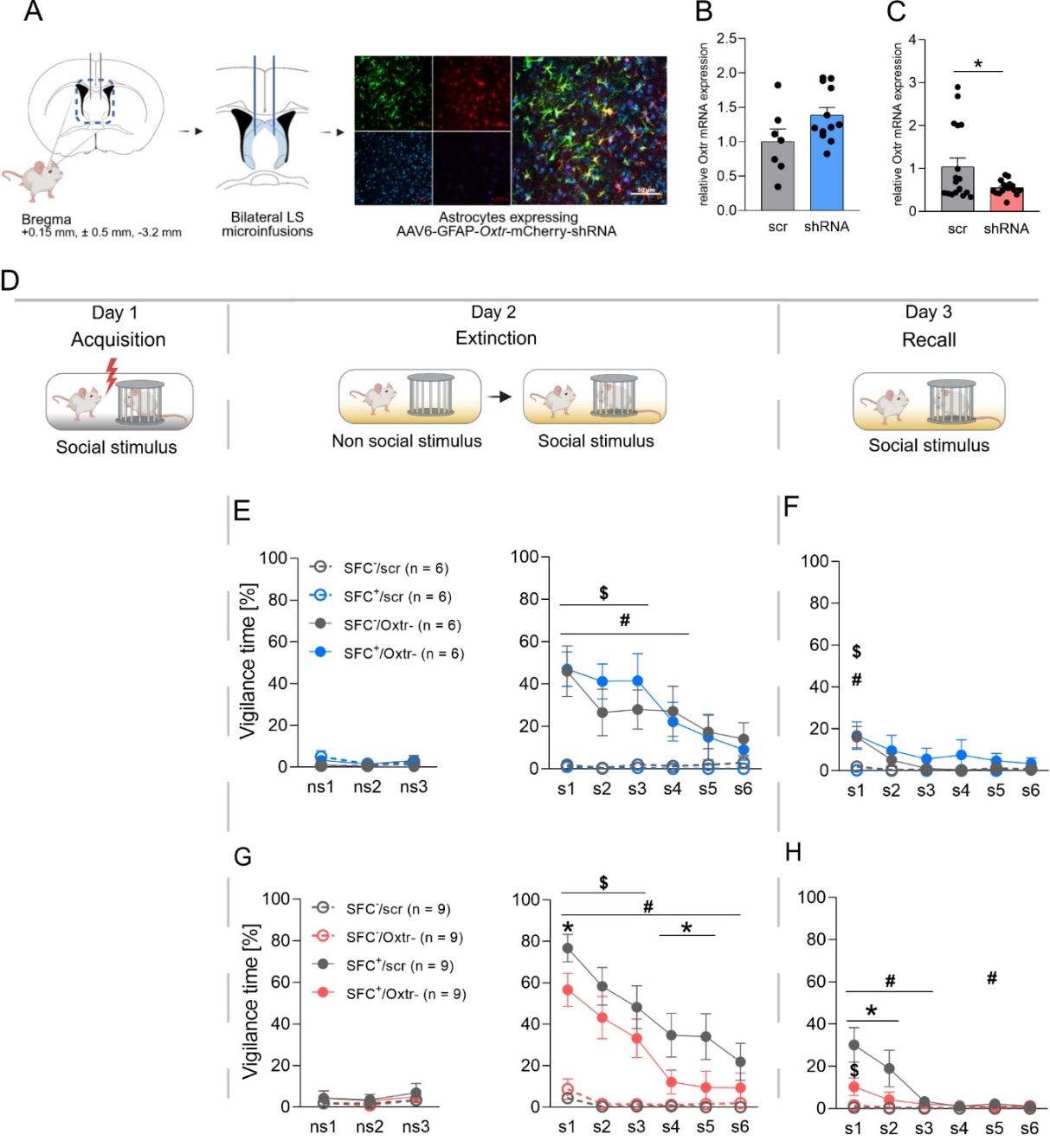

**Figure S6.** **Effects of OXTR mRNA knockdown (OXTR^-^) in caudal lateral septum (LSc) astrocytes on vigilance behavior during social fear extinction (E, G), and recall (F, H) in male (blue) and female (red) mice. A, D.** Schematic presentation of the surgical and experimental protocol. **B, C.** qPCR data showing Oxtr mRNA levels in the LSc of male (B) and female (C) mice. The LSc of male and female mice were infused with an AAV expressing an shRNA targeting OXTR mRNA (OXTR^-^) or an AAV expressing a scrambled control (scr) three weeks prior to the SFC paradigm. **E, G.** Percentage of investigation time of the three non-social (ns1 - ns3; left) and six social (s1 – s6; right) stimuli during social fear extinction (day 2) in male and female mice. **F, H.** Percentage of investigation time of the six social stimuli (s1 – s6) during recall (day 3) in male and female mice. Data represent means ± SEM. * p ≤ 0.05 SFC^+^/OXTR^-^ vs SFC^+^/Scr (G), # p ≤ 0.05 SFC^+^/Scr vs SFC^-^/Scr (E, F, G, H), $ p ≤ 0.05 SFC^+^/OXTR^-^ vs SFC^-^/OXTR^--^ (E, F, G). For detailed statistics, see Supplementary Tables T12 (1-7) and T13 (1-7).
